# Effects of insulin supplementation on senescence-associated markers and cellular phenotype of human dental pulp stem cell spheroids during long-term serum-free culture

**DOI:** 10.64898/2026.11.17.767347

**Authors:** Yudha Tirtadani, Elshimaa Mohamedali Ahmed Mohamed, Annisa Khumaira, Fuad Gandhi Torizal

## Abstract

**Background:** In vitro expansion of human dental pulp stem cells (DPSCs) in serum-free, three-dimensional suspension culture is important for scalable, clinically relevant manufacturing. However, this approach is limited by reduced proliferative capacity and passage-dependent senescence accumulation. In this study, we evaluated whether insulin supplementation could address these limitations while maintaining mesenchymal immunophenotype and stemness-related gene expression across serial passages in a downscale model as a reference for further DPSCs upscaling.

**Methods:** DPSCs were expanded as spheroids for five consecutive 8-day passages (40 days total) in serum-free medium (SF), serum-free medium supplemented with insulin (SF+I), or medium containing 10% fetal bovine serum (FBS). Spheroid diameter, cell-equivalent yield, and fold expansion per passage were quantified. qPCR analysis of proliferation-associated (Ki67, PCNA), senescence-associated (p16, p21), and stemness-associated (OCT4, SOX2, NANOG, KLF4) transcripts, SA-β-galactosidase staining, and flow-cytometric analysis of MSC-associated surface markers were performed.

**Results:** SF+I increased per-passage fold expansion relative to SF and produced expansion values that were comparable from those in FBS (22.3 fold vs 25.0 fold) with larger spheroid diameters. SF showed progressive upregulation of senescence marker p16 and p21 across passages (2.7-fold increase from passage 1 to passage 5), whereas SF+I maintained lower expression of both markers and showed markedly less SA-β-galactosidase positivity at passage 5, compared with SF-based medium (13.19% vs 46.81%). KLF4 and NANOG were better retained in Sf and SF+I under serum-free conditions than under FBS, although the statistical differences were marker-specific. SF and SF+I also retained high positivity for CD105, CD90, and CD73 (>90%), whereas FBS showed substantial loss of these MSC-associated markers (CD90: 31.6%; CD73: 28.4%).

**Conclusions:** Insulin supplementation supported long-term serum-free 3D DPSC expansion and was associated with lower senescence-associated readouts. Serum-free conditions were also associated with better preservation of the measured MSC-associated surface-marker profile and selected stemness-associated transcripts than FBS conditions. These findings warrant further multi-donor and functional validation of insulin-supplemented serum-free media for DPSC manufacturing.

## 1. INTRODUCTION

Human dental pulp stem cells (DPSCs) are mesenchymal stromal cells isolated from the dental pulp of permanent teeth (Shi et al., 2020). Teeth removed during routine dental care can provide an accessible source of cells, although donor and tissue variability remains relevant (Shi et al., 2020; Yamada et al., 2019). DPSCs exhibit proliferative capacity and multilineage differentiation in vitro (Al Madhoun et al., 2021; Yoshida et al., 2020; Luke et al., 2020; Yuan et al., 2022) and have been investigated for regenerative applications (Luke et al., 2020). Expansion is required to obtain sufficient cell numbers for research or potential applications, but serial passage can alter proliferation, differentiation, and senescence-associated features (Meng et al., 2022; Wang et al., 2018; Xu et al., 2024). Therefore, preserving cellular characteristics during expansion is an important experimental question question (Bakopoulou et al., 2017; Qu et al., 2020).

Three-dimensional spheroid culture alters cell–cell and cell–matrix interactions and has been studied as an alternative to monolayer culture. Human dental pulp cells can form spheroids in serum-free medium and persist through extended culture, whereas DPSC spheroid studies have reported differences in stemness-associated expression and differentiation compared with monolayer cultures (Xiao and Tsutsui, 2013; Labedz-Maslowska et al., 2020; Chan et al., 2021; Raik et al., 2023; Yen et al., 2023). Serum-free DPSC formulations have also yielded outcomes that vary by medium and assay, including differences in proliferation and surface-marker expression relative to serum-containing conditions (Abdel Moniem et al., 2019; Qu et al., 2019). Avoiding animal serum may reduce exposure to undefined xenogeneic components, but serum-free culture does not necessarily improve every measure of cell quality (Suchánek et al., 2013; Qu et al., 2020).

Insulin is a biologically plausible supplement to test in serum-free media because insulin receptor signaling can regulate cellular metabolism, growth, and survival in a context-dependent manner (Boucher et al., 2014; Kwon and Pessin, 2020; Solinas and Becattini, 2022). In human DPSCs, insulin has been reported to influence proliferation, osteogenic differentiation, and bone-forming outcomes, with pathway analyses implicating insulin/insulin-like growth factor signaling through PI3K, AKT, and mTOR (E et al., 2024). These findings provide a rationale for testing insulin during DPSC expansion but do not establish its effects during long-term serum-free spheroid culture. The present study therefore compared insulin-supplemented serum-free medium with serum-free medium alone and an FBS-containing condition across serial passages.

This downscaled study examined whether insulin supplementation was associated with differences in expansion, senescence-associated readouts, surface-marker expression, and selected transcripts during long-term serum-free three-dimensional culture of human DPSC spheroids. Spheroid morphology, cell-equivalent yield, and per-passage fold expansion were evaluated alongside MSC-associated surface markers, stemness-associated transcripts, and senescence-associated measurements.

### 2. MATERIALS AND METHODS

### 2.1. Maintenance and Spheroid Formation

DPSCs (Lonza Bioscience, Basel, Switzerland) were initially expanded in monolayer culture using Dental Pulp Stem Cell BulletKit^™^ Medium (Lonza Bioscience), with nutrients replaced every other day. To initiate the transition to three-dimensional (3D) aggregates, confluent monolayers were enzymatically detached with trypsin–EDTA (Thermo Fisher Scientific, MA, USA) to obtain a homogeneous single-cell suspension. Spheroid formation was induced by inoculating approximately 10^6^ cells into 4 mL of medium in ultra-low-attachment 6-well vessels (Corning, NY, USA). Cultures were maintained under orbital agitation at 90 rpm at 37 °C in a controlled 5% O2 atmosphere for an initial 24-hour aggregation phase. Subsequently, mature spheroids were transitioned into specific experimental conditions using α-MEM (Lonza Bioscience) based serum-free media (with or without insulin supplementation) or standard FBS-containing controls. This 3D spheroid suspension cultivation spanned five serial passages, each lasting 8 days, with media refreshed every 48 hours to maintain metabolic stability.

### 2.2. Quantification of Cell Populations

Total cell yield was estimated by fluorometric quantification of double-stranded DNA (dsDNA) with the Quant-iT PicoGreen assay (Thermo Fisher Scientific). After a PBS wash, spheroids were chemically lysed in a specialized buffer containing 0.02% SDS (Thermo Fisher Scientific) and 0.2 mg/mL proteinase K (Thermo Fisher Scientific) at 37 °C. The resulting lysates were incubated with PicoGreen dsDNA reagent in the dark for 5–10 minutes. Fluorescence intensity was measured with a Fluoroskan Ascent^™^ FL microplate reader (Thermo Fisher Scientific). Final cell counts were derived from a standard curve established using predefined DPSC concentrations processed under identical parameters.

### 2.3. Morphometric Assessment

To monitor spheroid population growth, images were acquired using an Olympus BX51 light microscope (Evident Scientific, Tokyo, Japan). The mean diameter of the spheroids was calculated using the Fiji/ImageJ platform (v1.54).

### 2.4. Gene expression analysis

Total cellular RNA was isolated using TRIzol^™^ reagent (Life Technologies), followed by cDNA synthesis using SuperScript^™^ IV VILO^™^ Master Mix (Thermo Fisher Scientific). Quantitative real-time PCR (qPCR) was conducted on the PikoReal Real-Time PCR System using PowerUp^™^ SYBR^™^ Green Master Mix, adhering to the manufacturer’s thermal cycling protocols. The primer sequence used in this study is listed in Supplementary table 1.

### 2.6. Immunophenotypic Characterization

For surface marker analysis, the 3D structures were first dissociated into a single-cell suspension by incubating with Accutase (STEMCELL Technologies, Vancouver, Canada) for 30 minutes at 37 °C, followed by mechanical filtration through a 40 µm strainer (Corning, NY, USA). To minimize non-specific binding, 106 cells were incubated with a 2% BSA blocking solution in PBS for 30 minutes. The cells were then incubated sequentially with primary and secondary antibodies at room temperature in the dark for 30 minutes each. Nuclear counterstaining was performed with DAPI (1:1000; Dojindo, Kumamoto, Japan). High-throughput data acquisition was performed on a BD FACS Canto II system (BD Biosciences, NJ, USA). The antibodies used in this study are listed in Supplementary table 2.

### 2.6. Assessment of Senescence-Associated β-Galactosidase Activity

To identify senescent cell populations within the 3D structures, β-galactosidase activity was assessed using a Senescence-associated β-galactosidase (SA-β-Gal) kit (Cell Signaling Technology, MA, USA) adapted for cryosectioned aggregates. MSC spheroids were harvested and stabilized for 10 minutes at room temperature with the kit-supplied fixative (containing glutaraldehyde, methanol, and formaldehyde). After fixation, the microtissues were embedded in Tissue-Tek O.C.T. compound (Sakura Finetek, Torrance, CA, USA) and cryopreserved at _−_80°C until hardened. For histological analysis, 10-µm sections were obtained with a cryostat (Leica Biosystems, Nussloch, Germany) and mounted on glass slides. SA-β-Gal staining was performed directly on slide-mounted cryosections by incubating them overnight at 37°C (without CO_2_) until the blue chromogenic signal was detectable under light microscopy. Following the enzymatic reaction, nuclei were counterstained with Mayer’s hematoxylin (Sigma-Aldrich, MO, USA) for 5 minutes at room temperature. Slides were air-dried and protected with an aqueous mounting medium (Dako, CA, USA). Microscopic evaluation was performed using an Olympus BX51 system, and resulting images were processed and analyzed with FIJI/ImageJ.

## 3. RESULTS

### 3.1. 3D suspension culture model and experimental design

The DPSC spheroids were maintained for five consecutive eight-day culture passages (passage 1–5; total, 40 days) under three conditions: serum-free medium (SF), serum-free medium supplemented with insulin (SF+I), and medium containing 10% fetal bovine serum (FBS) (Fig. 1). Spheroid formation was observed during the initial aggregation stage (passage 0). Spheroid morphology, expansion-related measurements, and proliferation- and senescence-associated transcript levels were evaluated during serial culture. MSC-associated surface-marker positivity was assessed at the end of passage 5.

**Figure 1.**
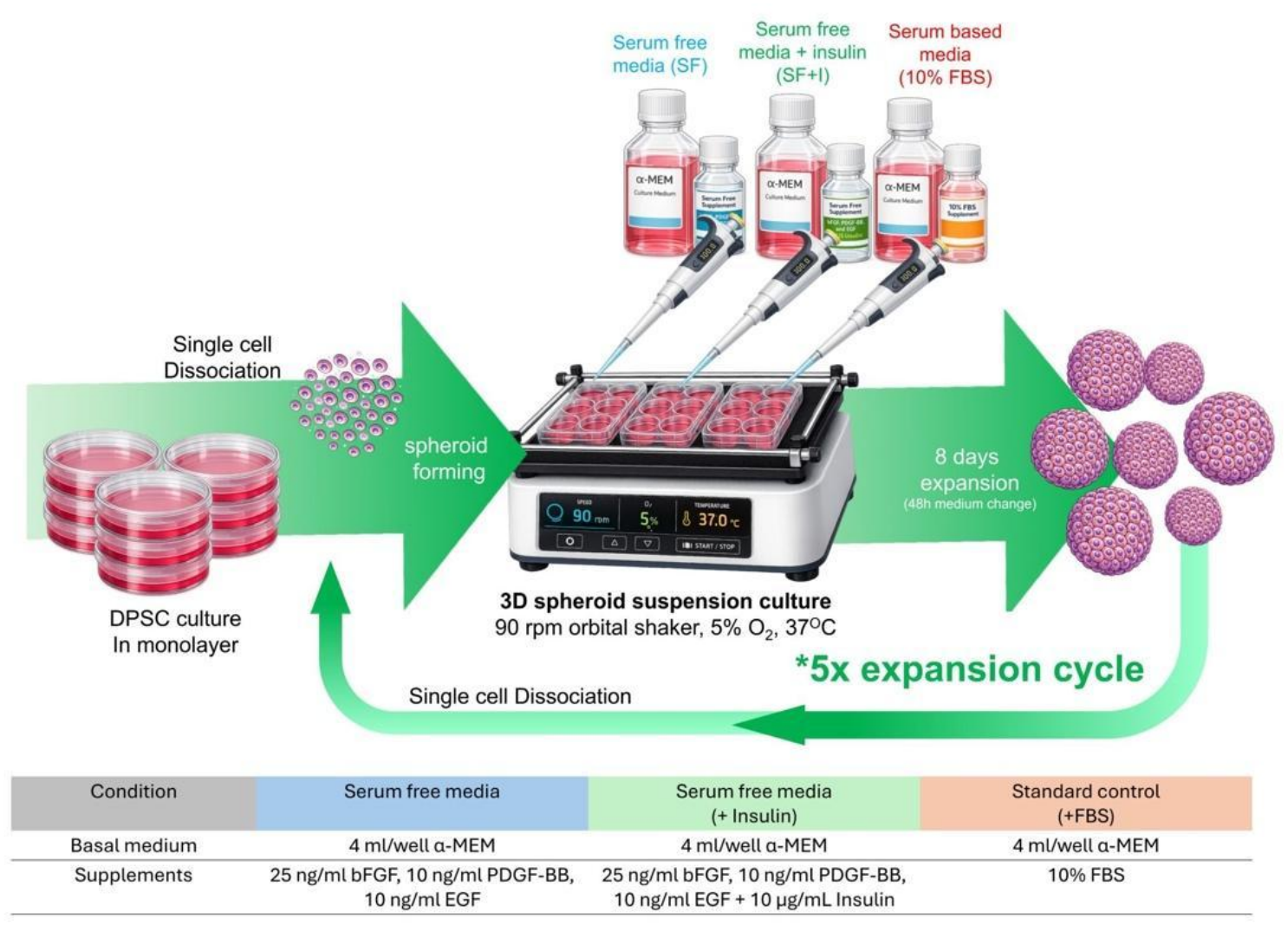
Experimental design for serial three-dimensional spheroid expansion of human dental pulp stem cells under serum-free, insulin-supplemented, and serum-containing conditions.

### 3.2. Insulin supplementation restores proliferative capacity during long-term serum-free expansion

Human dental pulp stem cells (DPSCs) formed compact, spherical aggregates under all three culture conditions, including serum-free medium (SF), insulin-supplemented serum-free medium (SF+I), and 10% fetal bovine serum (FBS), and maintained their spherical morphology throughout five consecutive passages (Fig. 2A). Morphometric analysis revealed differences in spheroid diameter across the culture conditions during serial expansion (Fig. 2B). At the initial aggregation stage (P0), mean spheroid diameters were similar across all three groups. From passage 1 (P1) onward, FBS generally produced the largest spheroids, followed by SF+I and SF, with this pattern maintained through passage 5 (P5). At P5, mean spheroid diameters were 263.65 ± 75.00 µm in SF, 350.64 ± 106.19 µm in SF+I, and 401.84 ± 112.73 µm in FBS. These observations indicate that insulin supplementation was associated with greater spheroid enlargement relative to SF alone during prolonged suspension culture.

**Figure 2.**
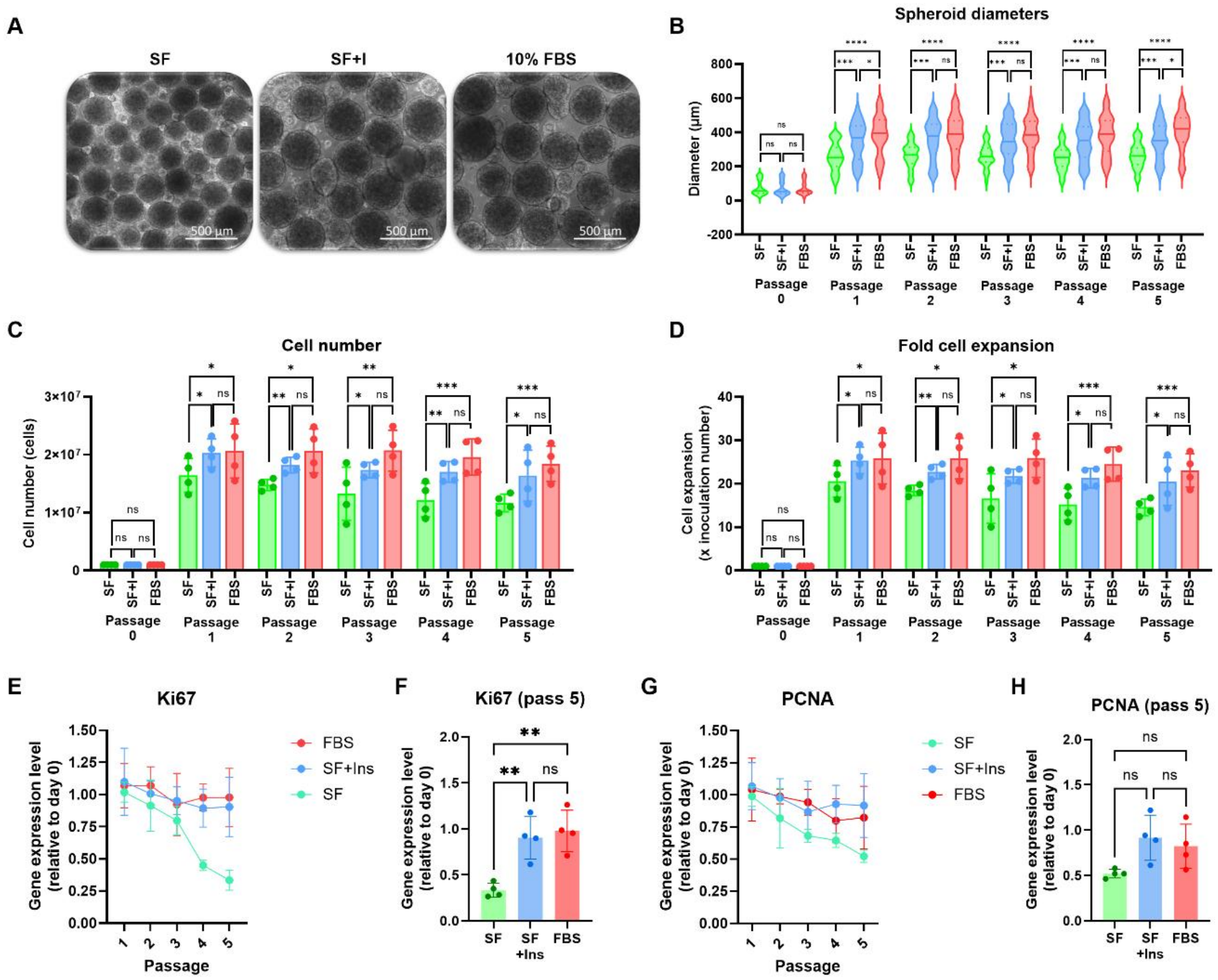
Insulin supplementation enhances proliferative capacity and supports sustained expansion of human dental pulp stem cells during long-term suspension culture. (A) Representative bright-field images of DPSC spheroids under the three culture conditions at passage 5. (B) Semi-quantitative image analysis of spheroid diameter across passages 0–5. (C) Total cell-equivalent yield per passage, estimated by PicoGreen double-stranded DNA quantification. (D) Fold expansion per passage across five consecutive passages. (E) Relative expression of the proliferation-associated marker Ki67 (MKI67) across passages 1–5 and (F) comparison of Ki67 expression at passage 5. (G) Relative expression of the proliferation-associated marker PCNA across passages 1–5 and (H) comparison of PCNA expression at passage 5. Gene expression was quantified using quantitative real-time PCR (qPCR). Data were obtained from three biologically independent replicates (n = 3). Statistical significance was assessed by one-way ANOVA followed by Tukey’s multiple-comparison test. Significance levels: ns, not significant; *P < 0.05; **P < 0.01; ***P < 0.001; and ****P < 0.0001.

Differences in spheroid size were accompanied by corresponding trends in total cell-equivalent yield and expansion capacity (Fig. 2C,D). DNA-based quantification showed that SF+I consistently produced higher mean cell-equivalent yields than SF across five consecutive passages, whereas FBS generally produced the highest yields (Fig. 2C). One-way ANOVA followed by Tukey’s multiple-comparison test revealed significantly higher cell-equivalent yields in FBS than in SF at passages 2–5. Although SF+I exhibited higher mean cell-equivalent yields than SF, these differences did not reach statistical significance at any passage. Similarly, no significant differences were detected between SF+I and FBS throughout the expansion period.

A comparable pattern was observed in per-passage fold expansion (Fig. 2D). The SF condition showed a progressive decline in mean fold expansion during serial culture, whereas SF+I and FBS maintained higher expansion values. At passage 5, mean fold expansion was 14.58 ± 1.90 in SF, 20.49 ± 5.52 in SF+I, and 23.05 ± 3.82 in FBS. One-way ANOVA with Tukey’s multiple-comparison test identified significantly higher fold expansion in FBS than in SF at passages 2–5. Although SF+I consistently showed higher mean fold expansion than SF, the differences did not reach statistical significance at any passage. Likewise, no significant differences were observed between SF+I and FBS throughout the five passages.

To further examine proliferation-associated transcriptional changes, the expression of Ki67 (MKI67) and proliferating cell nuclear antigen (PCNA) was evaluated across consecutive passages (Fig. 2E–H). Ki67 expression declined progressively in SF, particularly in later expansion cycles, whereas SF+I and FBS maintained higher transcript levels (Fig. 2E). This difference was also evident in the endpoint comparison at passage 5 (Fig. 2F). PCNA expression showed a more gradual passage-dependent decline, with lower transcript levels in SF than in SF+I and FBS in later passages (Fig. 2G). The corresponding passage 5 comparison further highlighted differences in PCNA expression among the culture conditions (Fig. 2H). These transcriptional patterns were generally consistent with the observed trends in cell-equivalent yield and fold expansion.

Collectively, these findings demonstrate that insulin supplementation was associated with larger spheroids, higher mean cell-equivalent yield, and greater mean fold expansion compared with serum-free medium alone during long-term three-dimensional suspension culture. Although the differences in cell-equivalent yield and fold expansion between SF and SF+I did not reach statistical significance in the passage-specific comparisons, the consistent numerical trends and proliferation-associated transcriptional profiles suggest that insulin supplementation may help sustain DPSC expansion under serum-free conditions.

### 3.3. Serum-free culture preserves mesenchymal immunophenotype independent of insulin supplementation

We evaluated the mesenchymal stromal cell (MSC)-associated immunophenotype of human dental pulp stem cells (DPSCs) by flow cytometry after five consecutive passages under serum-free (SF), insulin-supplemented serum-free (SF+I), and 10% fetal bovine serum (FBS) culture conditions (Fig. 3A–C). The analysis included the positive MSC-associated surface markers CD105, CD90, and CD73, together with a combined panel of negative lineage markers comprising CD19, CD11b, CD34, and CD45.

**Figure 3.**
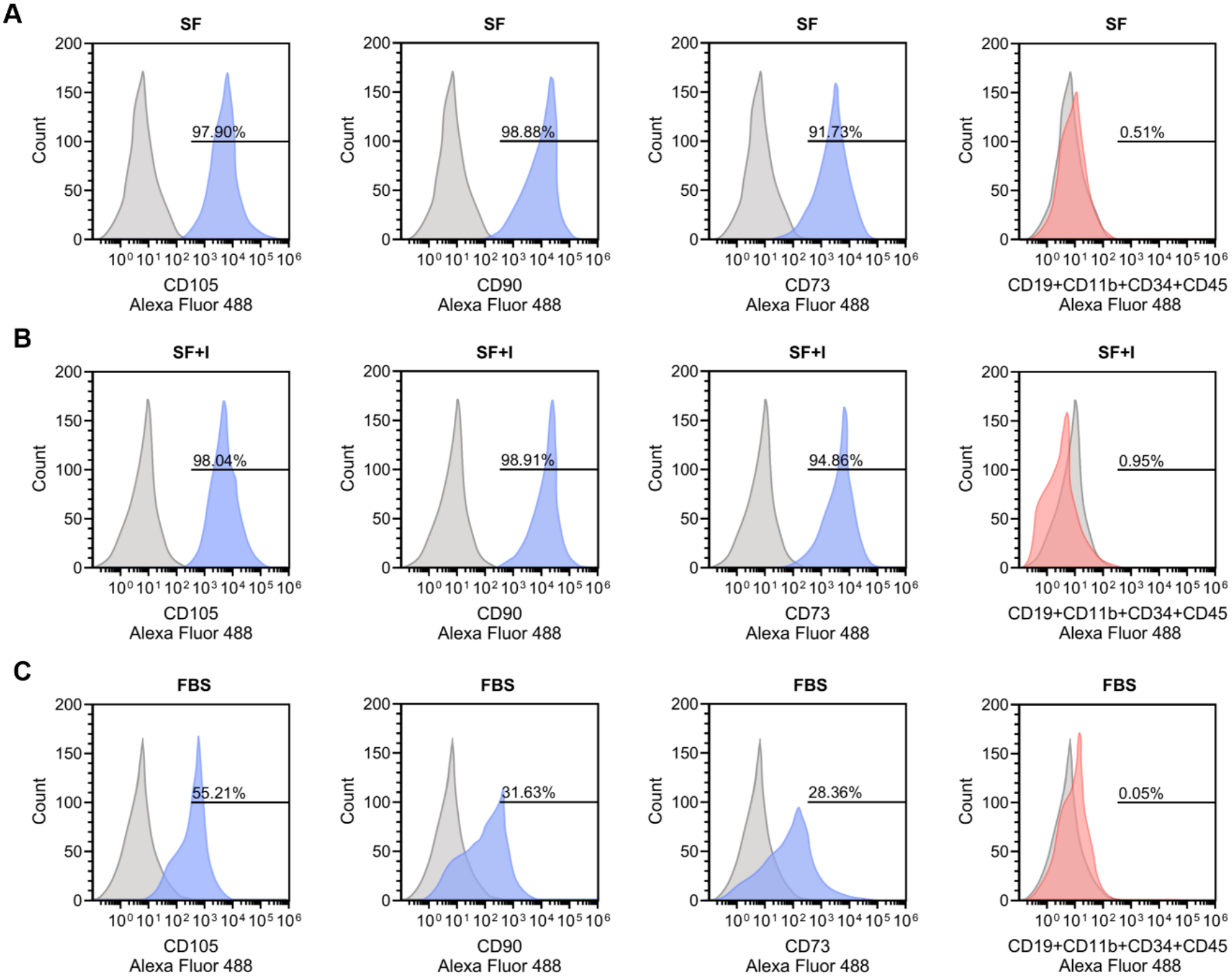
Serum-free culture conditions maintain mesenchymal stromal cell-associated surface marker expression during long-term three-dimensional suspension culture of human dental pulp stem cells. Flow cytometric analysis of mesenchymal stromal cell (MSC)-associated surface markers in human dental pulp stem cells (DPSCs) cultured under serum-free (SF), insulin-supplemented serum-free (SF+I), and 10% fetal bovine serum (FBS) conditions following five consecutive passages. Representative flow cytometry histograms show the expression of MSC-positive surface markers CD105, CD90, and CD73, and a combined panel of negative lineage markers (CD19, CD11b, CD34, and CD45) under (A) SF, (B) SF+I, and (C) FBS conditions. The percentages of marker-positive cells are indicated within each histogram. Gray histograms represent unstained controls, while colored histograms represent antibody-stained cell populations.

At passage 5 (P5), both serum-free conditions exhibited high positivity for the measured MSC-associated surface markers (Fig. 3A,B). In SF, the proportions of CD105, CD90, and CD73 positive cells were 97.9%, 98.9%, and 91.7%, respectively, while the corresponding values in SF+I were 98.0%, 98.9%, and 94.9%. Positivity for the combined negative lineage-marker panel remained below 1% in both conditions, with values of 0.51% and 0.95% in SF and SF+I, respectively. These findings indicate that the two serum-free formulations exhibited similar MSC-associated surface-marker profiles at the final passage, irrespective of insulin supplementation.

In contrast, DPSCs expanded under FBS-containing conditions exhibited markedly lower proportions of cells expressing the positive MSC-associated surface markers at P5 (Fig. 3C). CD105, CD90, and CD73 positivity was 55.2%, 31.6%, and 28.4%, respectively, whereas positivity for the combined negative lineage-marker panel remained low (0.05%). Compared with both serum-free conditions, the FBS group therefore exhibited a lower proportion of cells expressing the measured positive MSC-associated markers, despite maintaining low expression of the negative lineage-marker panel.

These observations demonstrate that SF and SF+I were associated with higher MSC-associated surface-marker positivity than FBS after long-term three-dimensional suspension culture. The similar immunophenotypic profiles observed in SF and SF+I suggest that insulin supplementation was not associated with substantial changes in the measured surface-marker profile at the final passage. These findings indicate that the serum-free culture environment was associated with greater retention of the investigated MSC-associated surface markers, although the endpoint analysis alone does not establish their stability throughout the preceding passages.

### 3.4. Serum-free conditions preserve stemness-associated transcriptional programs independently of insulin supplementation

The expression of four stemness-associated transcription factors, OCT4, SOX2, NANOG, and KLF4, was evaluated across five consecutive passages to assess how culture conditions affect their transcriptional profiles during long-term three-dimensional DPSC expansion (Fig. 4A–H). Although all three culture conditions exhibited passage-dependent changes in gene expression, the magnitude and pattern of these changes varied across the markers.

**Figure 4.**
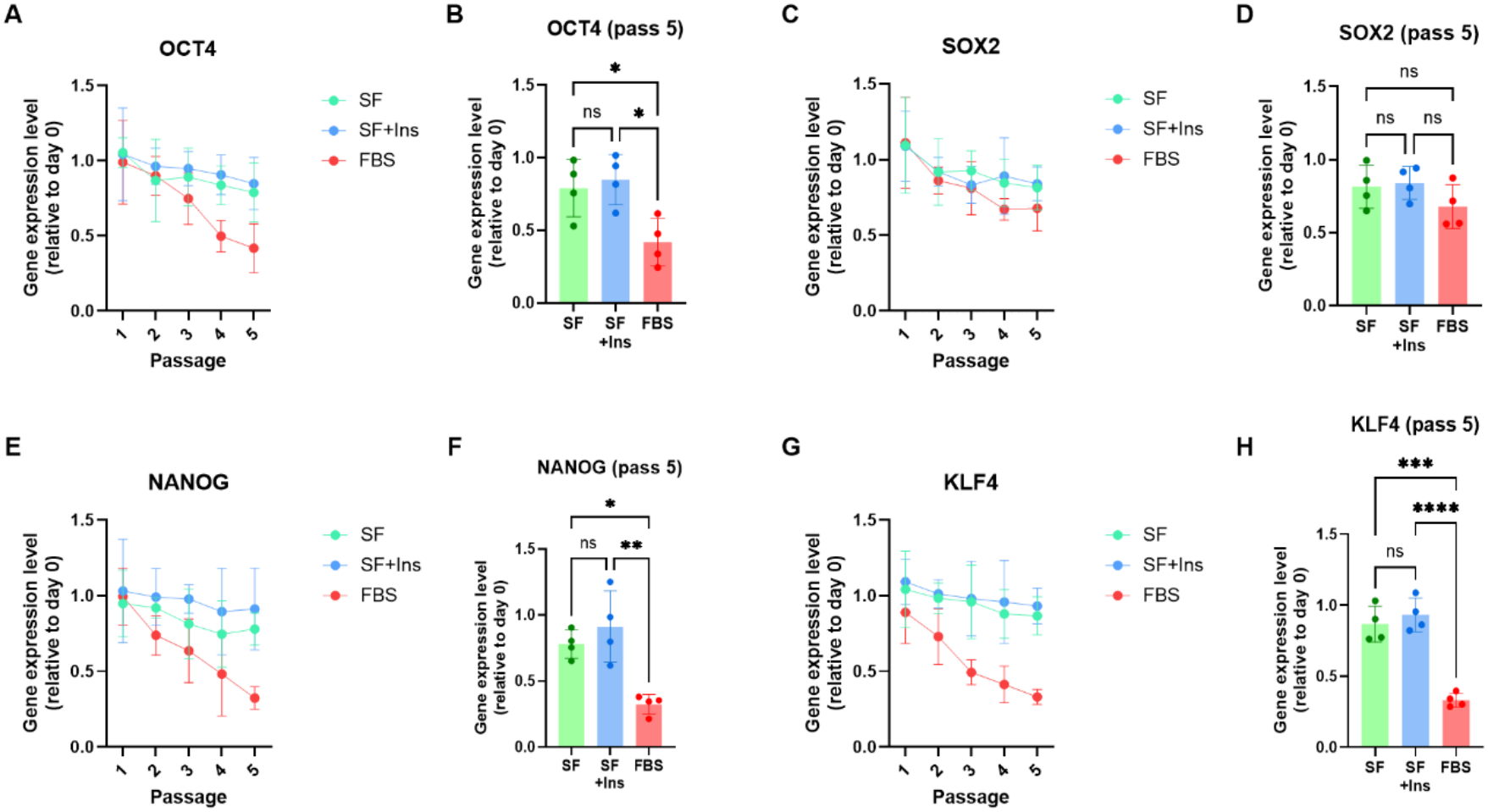
Expression of stemness-associated transcription factors in human dental pulp stem cells during long-term three-dimensional suspension culture. Quantitative real-time PCR (qPCR) analysis of stemness-associated transcription factors in human dental pulp stem cells (DPSCs) cultured under serum-free (SF), insulin-supplemented serum-free (SF+I), and 10% fetal bovine serum (FBS) conditions across five consecutive passages. (A) OCT4 expression across passages 1–5 and (B) comparison at passage 5. (C) SOX2 expression across passages 1–5 and (D) comparison at passage 5. (E) NANOG expression across passages 1–5 and (F) comparison at passage 5. (G) KLF4 expression across passages 1–5 and (H) comparison at passage 5. Data were obtained from at least three biologically independent replicates (n = 3). Statistical significance was assessed by two-way ANOVA with Tukey’s multiple-comparison test. Significance levels: ns, not significant; *P < 0.05; **P < 0.01; ***P < 0.001; and ****P < 0.0001.

OCT4 expression tended to decline during serial expansion, with a more pronounced reduction under FBS-containing conditions than under SF and SF+I, particularly in later passages (Fig. 4A). A significant overall effect of culture condition was detected, with significantly lower OCT4 expression in FBS than both in SF and SF+I at passage 5. In contrast, SOX2 expression showed a gradual passage-dependent decline across all three conditions (Fig. 4C), with no significant overall effect of culture condition. The endpoint comparison at passage 5 likewise showed no significant differences among SF, SF+I, and FBS (Fig. 4D).

NANOG expression showed a condition-dependent pattern during serial expansion, with a greater reduction in FBS than in serum-free conditions (Fig. 4E). At passage 5, NANOG expression was significantly lower in FBS than in SF+I, whereas the difference between FBS and SF did not reach statistical significance (Fig. 4F). These findings indicate greater retention of NANOG transcript expression under insulin-supplemented serum-free conditions relative to FBS at the final passage.

Among the investigated transcription factors, KLF4 showed the strongest overall condition-dependent difference, with changes becoming more apparent in later passages (Fig. 4G). At passage 5, KLF4 expression was lower in FBS than in SF and SF+I, whereas no significant difference was detected between SF and SF+I (Fig. 4H). These observations demonstrate greater retention of KLF4 transcript expression under both serum-free conditions compared with FBS at the end of serial expansion.

These overall results indicate that long-term culture conditions differentially influenced the expression of stemness-associated transcription factors in DPSCs. Serum-free conditions were associated with greater retention of selected transcripts, particularly KLF4 and NANOG, compared with FBS-containing medium. No significant differences between SF and SF+I were detected for the investigated markers in the reported comparisons, suggesting that the observed transcriptional differences were primarily associated with the serum-free culture environment rather than an independently demonstrated effect of insulin supplementation.

### 3.5. Insulin attenuates passage-dependent senescence marker accumulation in serum-free expansion

The expression of the senescence-associated cell-cycle inhibitors p16 (CDKN2A) and p21 (CDKN1A) was assessed across five consecutive passages to investigate how culture conditions affect senescence-associated transcriptional changes during long-term three-dimensional DPSC expansion (Fig. 5A–D). Both markers showed distinct expression patterns across the culture conditions, with consistently higher expression in serum-free medium (SF) than in insulin-supplemented serum-free medium (SF+I) and in fetal bovine serum (FBS).

**Figure 5.**
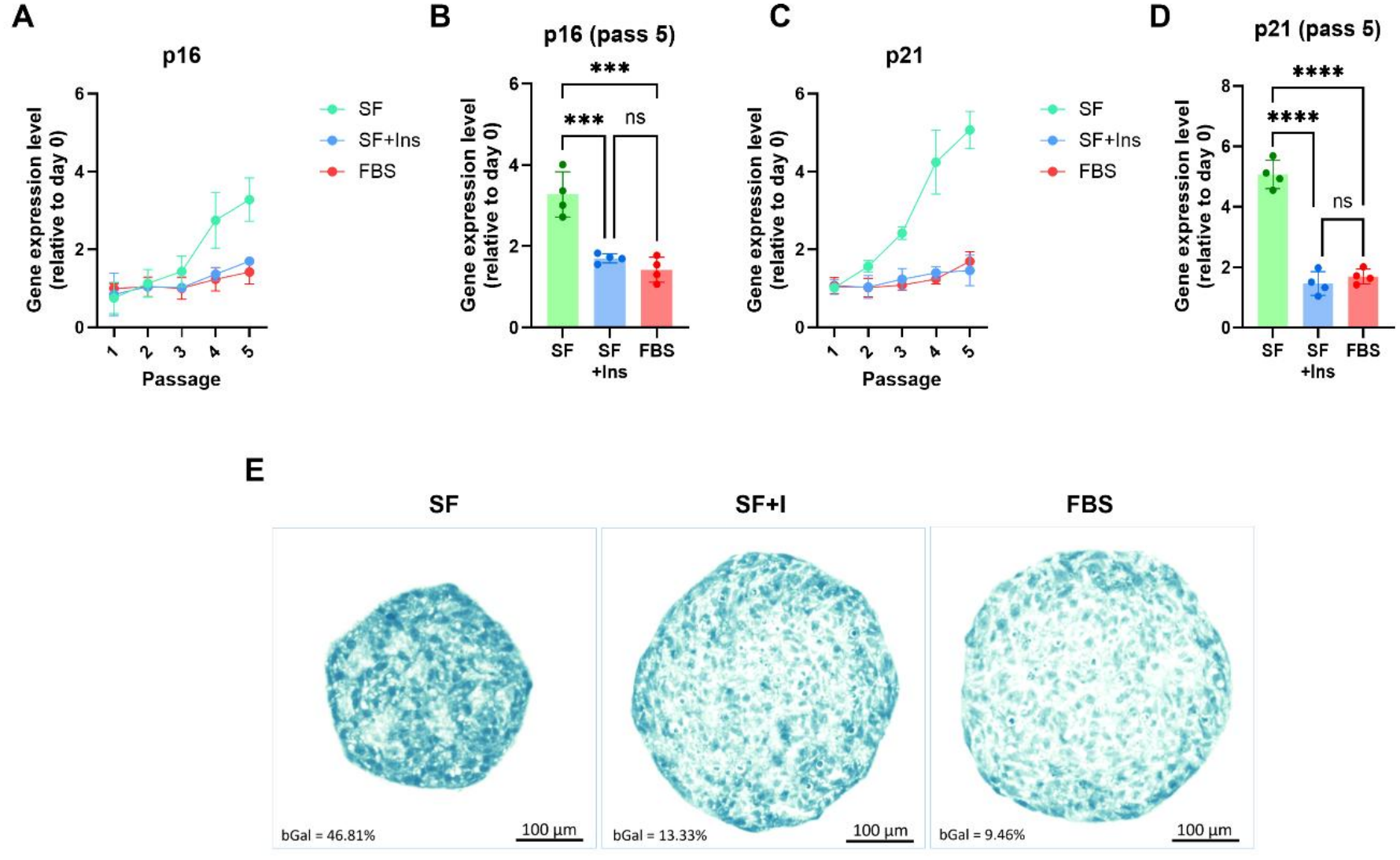
Insulin supplementation attenuates senescence-associated changes during long-term serum-free expansion of human dental pulp stem cells. Senescence-associated marker expression and senescence-associated β-galactosidase (SA-β-gal) activity were evaluated in human dental pulp stem cells (DPSCs) cultured under serum-free (SF), insulin-supplemented serum-free (SF+I), and 10% fetal bovine serum (FBS) conditions across five consecutive passages. (A) Relative expression of the senescence-associated marker p16 (CDKN2A) across passages 1–5 and (B) comparison of p16 expression level at passage 5. (C) Relative expression of p21 (CDKN1A) across passages 1–5 and (D) comparison of p21 at passage 5. Gene expression was quantified using quantitative real-time PCR (qPCR). (E) Representative histological images of SA-β-gal staining in DPSC spheroid cryosections at passage 5, showing SA-β-gal-positive cells (teal staining) and hematoxylin-counterstained nuclei. The percentages of SA-β-gal-positive cells were 46.81%, 13.19%, and 9.46% in SF, SF+I, and FBS, respectively. Data were obtained from three biologically independent experiments (n = 3). Statistical significance for gene-expression analyses was determined by one-way ANOVA followed by Tukey’s multiple comparisons test. Significance levels: ns, not significant; *P < 0.05; **P < 0.01; ***P < 0.001; and ****P < 0.0001.

p16 expression was consistently elevated in SF throughout serial expansion, whereas SF+I and FBS maintained lower levels (Fig. 5A). Statistical analysis revealed a significant overall effect of culture condition, with significantly higher p16 expression in SF than in SF+I and FBS across all five passages. We detected no significant differences between SF+I and FBS at any passage. This pattern was also evident at passage 5 (P5), where SF exhibited higher p16 expression than either SF+I or FBS (Fig. 5B).

A similar condition-dependent pattern was observed for p21, although its temporal expression profile differed from that of p16 (Fig. 5C). In SF, p21 expression increased progressively across consecutive passages, reaching approximately 2.7-fold the level at passage 1 (P1) by P5. In contrast, SF+I showed a less pronounced passage-dependent increase, while FBS maintained relatively low expression throughout the expansion period. Statistical analysis revealed a significant overall effect of culture condition, with no significant differences between SF+I and FBS in the reported passage-specific comparisons. At passage 5, p21 expression remained higher in SF than in the other culture conditions (Fig. 5D).

To complement the transcriptional analyses, senescence-associated β-galactosidase (SA-β-gal) activity was assessed histochemically in cryosections of DPSC spheroids at P5 (Fig. 5E). Semi-quantitative image analysis showed that the proportion of SA-β-gal-positive cells was highest in SF (46.81%), followed by SF+I (13.19%) and FBS (9.46%). Compared with SF, insulin supplementation was associated with a substantially lower proportion of SA-β-gal-positive cells at the final passage. This descriptive pattern was consistent with the lower p16 and p21 transcript levels observed in SF+I relative to SF.

The detection of senescence markers indicates that insulin supplementation was associated with attenuation of senescence-associated changes during long-term, serum-free DPSC expansion. The lower expression of p16 and p21, together with the reduced proportion of SA-β-gal-positive cells, suggests that insulin supplementation may help limit the accumulation of senescence-associated features during serial three-dimensional suspension culture.

## 4. DISCUSSIONS

DPSCs are an accessible oral cell source with reported proliferative and multilineage differentiation capacity, but their characteristics can change during in vitro expansion (Wang et al., 2018; Yamada et al., 2019; Labedz-Maslowska et al., 2020; Torizal et al., 2022). Prior work has shown that human dental pulp cells can form spheroids under serum-free conditions and that DPSC spheroid culture can alter measured differentiation and stemness-associated outcomes (Xiao and Tsutsui, 2013; Chan et al., 2021; Raik et al., 2023). In the present 40-day comparison, insulin-supplemented serum-free culture was associated with numerically higher expansion and lower senescence-associated readouts than serum-free medium alone. Because the study used one established cell line and several expansion comparisons were not statistically significant, these findings should be interpreted as preliminary evidence specific to the tested culture conditions.

At the end of culture, mean fold expansion was higher in SF+I than in SF, but the difference did not reach statistical significance. Cell-equivalent yield comparisons showed the same limitation. The pattern is therefore suggestive rather than evidence that insulin restored proliferation. A separate study in human DPSCs reported insulin-associated effects on proliferation and osteogenic outcomes and implicated the IIS/PI3K/AKT/mTOR pathway (E et al., 2024), but its findings do not establish the mechanism in the present spheroid-expansion model. Insulin receptor signaling can affect growth and metabolism through several pathways (Boucher et al., 2014; Kwon and Pessin, 2020; Solinas and Becattini, 2022). Here, receptor activation, downstream signaling, nutrient use, and metabolic flux were not measured, so any mechanistic explanation remains a hypothesis.

In general, SF and SF+I showed greater positivity for measured MSC-associated surface markers than FBS. Similar SF and SF+I endpoint profiles do not indicate an independent insulin effect, and a single endpoint cannot establish maintenance across passages. Transcript results varied by marker. OCT4, KLF4, and NANOG differed in some comparisons, while SOX2 showed no significant condition effect. These transcripts should not be treated as stand-alone evidence of stem-cell identity or potency. In cultured adult human MSCs, NANOG expression has been associated with adaptation to in vitro growth and did not only identify stem or progenitor properties (Pierantozzi et al., 2011). The observed differences may relate to the culture conditions tested with insulin, but this study did not identify which components account for the FBS-versus-serum-free differences. Prior DPSC studies likewise show that serum-free outcomes vary with the formulation and assay used (Abdel Moniem et al., 2019; Qu et al., 2020; Suchánek et al., 2013).

The senescence findings provide a coherent explanation for the proliferative divergence between SF and SF+I that goes beyond a straightforward deficit in mitogenic input. Cellular senescence can be initiated by extrinsic and intrinsic stimuli, including telomere attrition, nutrient deprivation, and mitochondrial dysfunction, and its progression is governed by two principal regulatory axes, which are p53/p21 and p16/RB tumor suppressor pathways (Kumari and Jat, 2021). Within these pathways, p21 primarily initiates senescence-associated growth arrest, whereas p16 is required to maintain the arrested state once established (Itahana et al., 2001; Kumari and Jat, 2021). This distinction is relevant to the present findings, where p21 showed a progressive accumulation trajectory in SF cultures across passage, while p16 remained consistently elevated throughout, a pattern that aligns with a culture environment in which senescence is both continuously initiated and sustained in the absence of adequate growth support. Although no single senescence marker provides a fully reliable readout in isolation (Gorgoulis et al., 2019; Kumari and Jat, 2021), convergent evidence from p16 expression, p21 accumulation kinetics, and SA-β-galactosidase positivity across three distinct conditions strengthens the interpretation that the absence of insulin was associated with an increased senescence-associated profile that accumulated over time. Insulin may counteract this trajectory through AKT-mediated survival signaling, attenuation of FOXO-driven stress responses, or metabolic stabilization that reduces the burden of growth factor deprivation, though these pathways were not directly measured here and the mechanistic basis remains to be confirmed (Boucher et al., 2014; E et al., 2024; Kwon and Pessin, 2020). The present data indicate that insulin absence during serum-free expansion of an arrested population precludes assessment of functional fitness for downstream therapeutic use based on yield alone.

The findings are directly relevant to designing scalable DPSC manufacturing strategies. In cell therapy manufacturing, a culture condition that supports short-term proliferation but promotes phenotypic instability, senescence accumulation, or identity drift over extended passages is suboptimal, regardless of the cell numbers it generates (Torizal et al., 2019). Evaluating culture conditions across multiple serial passages, as done here, provides a more meaningful basis for manufacturing decisions than single-timepoint, short-duration assessments. These data suggest that insulin supplementation may support efficient expansion while preserving cellular characteristics over a clinically relevant expansion timeline. This benefit may be achievable in a suspension spheroid platform inherently compatible with scale-up bioprocessing (Torizal et al., 2026). Identifying medium components that simultaneously support yield and quality in defined, serum-free conditions is a prerequisite for reproducible and regulatory-compatible DPSC manufacturing, and insulin represents a biologically rational and practically accessible candidate for inclusion in such formulations.

Several limitations define the scope of the present findings and point toward necessary follow-up. First, this study used a single established DPSC line, and variability in insulin responsiveness, proliferation kinetics, senescence trajectory, and surface marker stability across donors may be substantial. Multi-donor validation is required before these findings can be generalized. Second, while the results are consistent with insulin’s known roles in PI3K/AKT-mediated growth and metabolic regulation, the study did not directly examine receptor activity, downstream signaling, or metabolic responses, so the proposed mechanisms remain inferential. Third, transcriptional and phenotypic maintenance alone does not confirm therapeutic potency. Functional differentiation assays are necessary to determine whether the quality attributes preserved under SF+I translate into retained biological activity relevant to regenerative applications.

## 5. CONCLUSIONS

In this small-scale study, insulin supplementation was associated with numerically higher expansion and lower senescence-associated readouts in serum-free DPSC spheroid culture. These findings support insulin supplementation as a promising strategy for serum-free DPSC expansion, warranting further validation of functional potency, reproducibility, and scalability before its application in standardized, clinically compatible, scalable DPSC manufacturing systems for regenerative medicine.

## Author Contributions

YT: investigation, formal analyses, manuscript writing. EMAM: investigation, formal analyses, manuscript writing. AK: review and editing. FGT: conceptualization, formal analyses, review and editing.

## Data Availability

The datasets analyzed during the current study are available from the corresponding author upon reasonable request.

## Declaration of AI-assisted technologies in the manuscript preparation process

During the preparation of this manuscript, the authors used Grammarly AI to improve grammar, clarity, and readability. The authors reviewed and edited the AI-assisted text and take full responsibility for the final content.

