## Supplementary material for "Effects of insulin supplementation on senescence-associated markers and cellular phenotype of human dental pulp stem cell spheroids during long-term serum-free culture"

**Supplementary table 1.** The primer sequences used in quantitative PCR (qPCR) assays for analyzing gene expression markers in this study

| Gene | Forward primer | Reverse primer |
| --- | --- | --- |
| Ki67 | GAAAGAGTGGCAACCTGCCTTC | GCACCAAGTTTTACTACATCTGCC |
| PCNA | CAAGTAATGTCGATAAAGAGGAGG | GTGTCACCGTTGAAGAGAGTGG |
| OCT3/4 | GACAGGGGGAGGGGAGGAGCTAGG | CTTCCCTCCAACCAGTTGCCCAAAC |
| SOX2 | GGGAAATGGGAGGGGTGCAAAAGAGG | TTGCGTGAGTGTGGATTGGTG |
| NANOG | AAAGAATCTTCACCTATGCC | GAAGGAAGAGGAGAGACAGT |
| KLF4 | CGAACCCACACAGGTGAGAA | GAGCGGGCGAATTTCCAT |
| p16 | CCTCGTGCTGATGCTACTGA | CATCATCATGACCTGGTCTTCT |
| p21 | GCTTCATGCCAGCTACTTCC | CCCTTCAAAGTGCCATCTGT |

**Supplementary Table 2.** The list of antibodies used for flow cytometry analysis.

| Protein target | Primary antibody/catalog number | Secondary antibody/catalog number |
| --- | --- | --- |
| CD90 | Rabbit Anti-human CD90 / Thy1 IgG (ab307736) | Goat Anti-Rabbit IgG H&L (Alexa Fluor® 488) (ab150077) |
| CD73 | Rabbit Anti-human CD73 IgG (ab317462) | Goat Anti-Rabbit IgG H&L (Alexa Fluor® 488) (ab150077) |
| CD105 | Rabbit Anti-human CD105 IgG (ab231774) | Goat Anti-Rabbit IgG H&L (Alexa Fluor® 488) (ab150077) |
| CD45 | Rabbit Anti-human CD45 IgG (ab40763) | Goat Anti-Rabbit IgG H&L (Alexa Fluor® 488) (ab150077) |
| CD34 | Rabbit Anti-human CD34 IgG (ab315802) | Goat Anti-Rabbit IgG H&L (Alexa Fluor® 488) (ab150077) |
| CD11b | Rabbit Anti-human CD11b IgG (ab133357) | Goat Anti-Rabbit IgG H&L (Alexa Fluor® 488) (ab150077) |
| CD19 | Mouse Anti-human CD19 IgG (ab136320) | Goat Anti-Mouse IgG H&L (Alexa Fluor® 488) (ab150113) |
